# The secreted protein signature of hydatid fluid from pulmonary cystic echinococcosis

**DOI:** 10.1101/2020.07.09.195701

**Authors:** Guilherme Brzoskowski dos Santos, Edileuza Danieli da Silva, Eduardo Shigueo Kitano, Maria Eduarda Battistella, Karina Mariante Monteiro, Jeferson Camargo de Lima, Henrique Bunselmeyer Ferreira, Solange Maria de Toledo Serrano, Arnaldo Zaha

**Author notes:** (G.B.S.); (E.D.S.); (M.E.B.) (K.M.M.); (J.C.L.); (H.B.F.) (E.S.K.); (S.M.T.S.). These authors contributed equally to this work (Guilherme Brzoskowski dos Santos, Edileuza Danieli da Silva).

## Abstract

The vast majority of cystic echinococcosis cases in Southern Brazil are caused by *Echinococcus granulosus* and *Echinococcus ortleppi*. Comparative proteomic studies of helminths have increased the knowledge about the molecular survival strategies adopted by parasites. Here, we surveyed the protein contents of the hydatid fluid compartment of *E. granulosus* and *E. ortleppi* pulmonary bovine cysts, in an attempt to compare their molecular arsenal in this host-parasite interface. Hydatid fluid samples from three isolates of each species were analyzed by trypsin digestion and mass spectrometry. We identified 280 proteins in *E. granulosus* and 251 proteins in *E. ortleppi*, highlighting a core of 52 proteins common to all samples of hydatid fluid. The *in silico* functional analysis revealed important molecular functions and processes active in pulmonary cystic echinococcosis. Some were more evident in one species, such as apoptosis in *E. ortleppi*, and cysteine protease activity in *E. granulosus*, while many molecular activities have been found in fluids of both species, such as proteolysis, development signaling and extracellular structures organization. The similar molecular tools employed by *E. granulosus* and *E. ortleppi* for their survival within the host are potential targets for new therapeutic approaches to deal with cystic echinococcosis and other larval cestodiases.

## 1. Introduction

Echinococcosis is a disease resulting from infection by flatworms of the *Echinococcus* genus. Depending on the species causing the infection, distinct features are noticed as a result of the differential development of the larval stage [1]. *Echinococcus granulosus* sensu stricto (s.s.; G1, G2 and G3 genotypes) and *E. ortleppi* are both etiological agents of cystic echinococcosis, in which the larval stage (metacestode) is characterized by the development of a unilocular, fluid-filled cyst (the hydatid cyst) in viscera from suitable intermediate hosts (mainly cattle and sheep). Humans can be accidental hosts and develop cystic echinococcosis as well. Due to their intermediate host preferences, *E. granulosus* s. s., from now on referred just as *E. granulosus*, and *E. ortleppi* were formerly known as *Echinococcus granulosus* sensu lato (s.l.) sheep and cattle strains, respectively [2,3]. After refinements in phylogenetic studies, *E. granulosus* s.l. cattle strain gained the status of species, *E. ortleppi* [3,4].

In the *E. granulosus* or *E. ortleppi* life-cycle [5], intermediate hosts get infected upon ingestion of eggs. Egg hatching releases oncospheres, which develop into hydatid cysts in viscera (mainly liver and lungs). A hydatid cyst is externally formed by an acellular, mucin-based laminated layer, and, internally, by a germinative layer. The germinative layer gives rise to brood capsules, where pre-adults (protoscoleces, PSCs) are produced by asexual reproduction. When PSCs are ingested by definitive hosts (canids, such as domestic dogs or wolves), they mature into adult worms within the small intestine, where they produce eggs that are released to the environment with host feces.

The hydatid cyst causes a chronic infection, as it can survive and grow for decades in the host, in most cases remaining fertile, with full capacity to generate PSCs [6]. To achieve this, the parasites adopt a wide repertoire of molecular strategies to both evade host defense mechanisms and acquire nutrients necessary for their development [7]. Such strategies allow parasite survival and development despite the chronic exposure to hostile environment provided by the host responses against infection. The liquid that fills the hydatid cyst, the hydatid fluid (HF), contains both parasite’s excretory-secretory (ES) products and host proteins, making it a rich component for the analysis of relevant molecules and interactions in a host-parasite interface [8,9].

Molecular characterization of the HF content is essential for a better understanding of the infection caused by *Echinococcus* spp., as well as for the discovery of new molecules with potential use in cystic echinococcosis diagnosis and treatment. Proteomic studies of helminth ES products have been particularly valuable for the identification of proteins involved in host-parasite relationship [10–12]. Prior *Echinococcus* spp. ES proteomic studies included the analysis of different components of *E. granulosus* cysts [8]; and comparisons among hydatid cyst fluid of *E. granulosus* cysts from different hosts (sheep, cattle and humans) [9]; and the comparison of HF from two different isolates of *Echinococcus multilocularis*, the etiological agent of alveolar echinococcosis [13]. Within the genus *Echinococcus*, interspecies comparisons in proteomic studies have been performed only between *E. granulosus* and *E. multilocularis* [14]. From all these studies, it became evident that analyses of the same species infecting different hosts, and different genotypes/species/strains infecting a common host, can provide valuable insights on molecular survival strategies adopted by parasites. The discovery of proteins shared by distinct species allows to outline conserved mechanisms involved in their interactions with the respective hosts. Furthermore, a species-specific set of proteins can provide molecular markers for parasite diagnosis.

So far, there were no proteomic comparative studies between two species of the *E. granulosus* s. l. complex. In this sense, *E. granulosus* and *E. ortleppi* constitute interesting subjects. Despite of its preference for ovine hosts, *E. granulosus* can also successfully infect, grow and asexually reproduce in bovine hosts, although with less efficiency than *E. ortleppi* [6,15]. While the fertility rate of *E. ortleppi* bovine cysts is very high (>90%), for *E. granulosus* it normally does not exceed 30% in this host species [2,16–18]. Moreover, *E. ortleppi* develops preferentially in lungs, whereas *E. granulosus* bovine cysts locate in both liver and lungs [19–22]. Therefore, *E. granulosus* and *E. ortleppi* bovine cysts offer the opportunity to comparatively assess two related species with different degrees of adaptation to a single host species.

Despite the above mentioned differences, hydatid cysts from *E. granulosus* and *E. ortleppi* are quite similar at macroscopic level. However, so far, no study has addressed their possible differences at the protein level, which may be related to the differential adaptation to survival and growth in a bovine host. Here, we generated protein profiles of HF from *E. granulosus* and *E. ortleppi* and compared the proteins identified to describe: i) the set of proteins shared between the two species and potentially involved in conserved mechanisms of parasite survival; and ii) the sets of distinct proteins that could indicate differences between these two species. The protein profiles generated showed a predominance of parasite proteins in the HF samples in comparison to the host proteins that infiltrate the cysts. Also, major molecular pathways acting in cattle pulmonary infections caused by *E. granulosus* and/or *E. ortleppi* were unveiled, helping to understand biologic aspects of the infection. The results presented in this work may assist the selection of potential targets in studies searching for new therapeutics, or disease markers capable of differentiating the two etiological agents.

## 2. Results

### 2.1. Protein Profiles from E. granulosus and E. ortleppi Hydatid Fluid Samples

A comparative proteomic survey was performed in an attempt to describe the HF protein components of *E. granulosus* and *E. ortleppi*, and to highlight differences and similarities between their protein repertoires. Because *E. ortleppi* develops predominantly in lungs [20,21], and to minimize differences in the protein profiles due to hydatid cyst location or the host species, we only used samples from pulmonary bovine infections. Three biological replicates, *i.e*., HF samples from individual fertile hydatid cysts, were analyzed for each species (EG1-3, for *E. granulosus*, and EO1-3, for *E. ortleppi*) and the numbers of identified proteins are summarized in Figure 1. To visualize the overall sample composition a heat map was performed with NSAF (Normalized spectral abundance factor) values of all identified proteins (Figure S1). The number of proteins identified vary among individual samples from each species. We were able to identify 207, 230 and 78 parasitic proteins in EG1, EG2 and EG3 HF samples, respectively, resulting in the overall identification of 280 *E. granulosus* unique proteins (Table S1A). In *E. ortleppi* HF samples, we identified 251 unique parasitic proteins, of which 194 were found in EO1, 224 in EO2 and 123 in EO3 (Table S1B). Overall, 214 proteins were shared between *E. granulosus* and *E. ortleppi*, 66 proteins were found exclusively in *E. granulosus*, and 37 proteins were found exclusively in *E. ortleppi*, totalizing 317 proteins. Proteins in the shared group did not show differences in abundance between *E. granulosus* and *E. ortleppi*, indicating that the two species may employ similar molecular strategies at the host-parasite interface (Figure S1 and Table S2).

**Figure 1.**
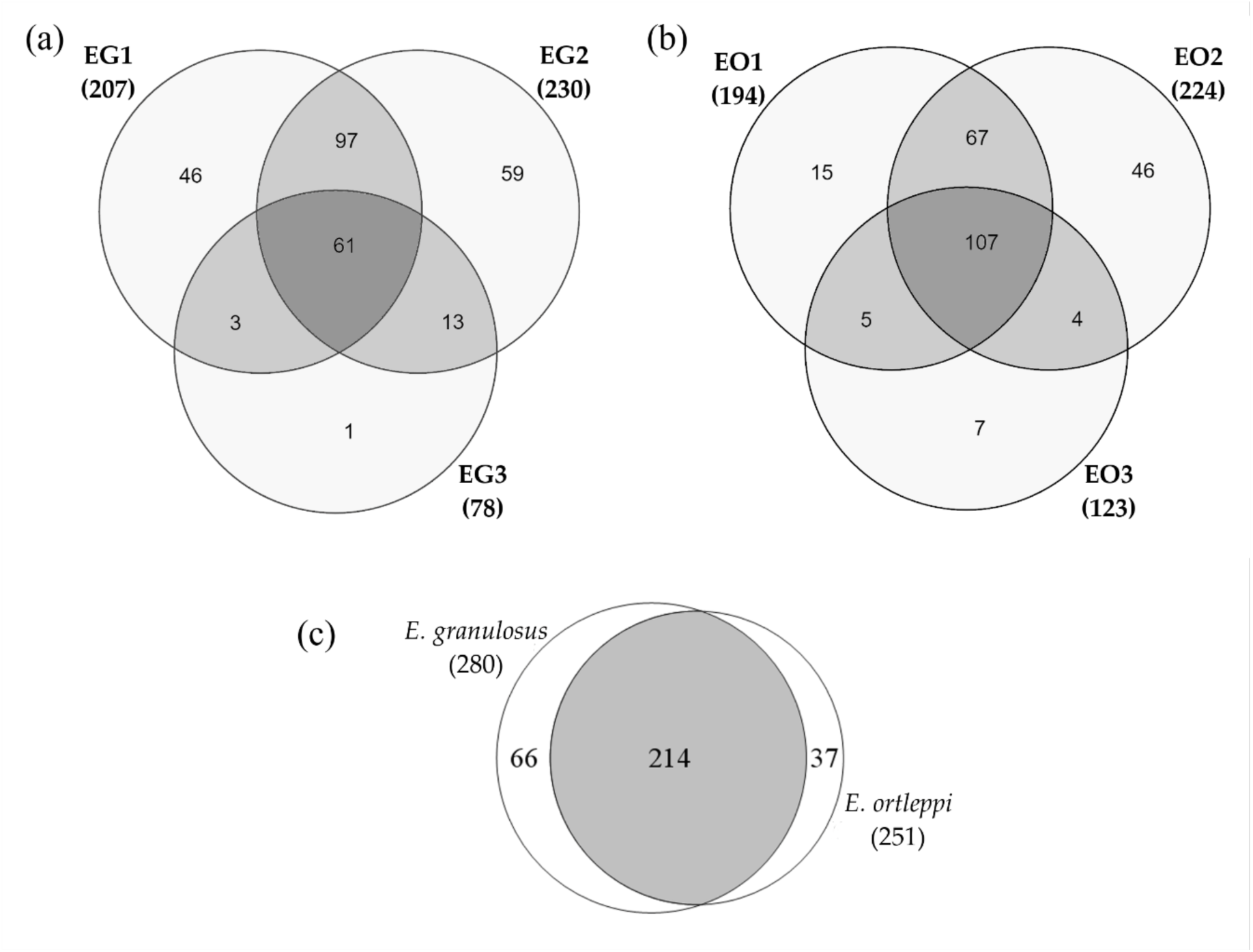
Parasitic proteins identified in HF samples from pulmonary cystic echinococcosis. Venn diagrams showing the number of proteins identified: (a) in *E. granulosus* biological replicates; (b) in *E. ortleppi* biological replicates; (c) in each species or shared between them. The overall numbers of proteins detected are indicated below the sample/species identification.

It is worth mentioning that a large group of proteins, 35 unique sequences, with unknown function was identified (Table S1). They were annotated as “expressed conserved protein”, “expressed protein” or “N/A (non-annotated)”. These proteins would be interesting study targets in order to elucidate their biological function. Some of these proteins with unknown function were identified in all the six samples, and some are among the most abundant proteins considering each species separately. Understanding their biological role will be an important contribution to the knowledge on *Echinococcus* spp biology.

As expected, host proteins were also identified in *E. granulosus* and *E. ortleppi* HF samples. However, host proteins were identified in smaller numbers in comparison to parasites’ proteins. Overall, 58 distinct *Bos taurus* proteins were identified, with 40 of them being identified in *E. granulosus* HF samples and 45 in *E. ortleppi* ones (Table S3). Variable numbers of bovine proteins were found in each biological sample, 12, 13 and 28 for EG1, EG2 and EG3, respectively, and 21, 11 and 37 for EO1, EO2 and EO3, respectively (Figure S2).

#### 2.1.1. Main Parasitic Proteins Identified in Hydatid Fluid Samples from *E. granulosus* and *E. ortleppi*

To highlight the most frequent parasitic proteins in HF, we selected those detected in at least two samples in one of the species, in a total of 217 proteins, from which 13 and 15 were detected exclusively in *E. granulosus* and *E. ortleppi* samples, respectively (Table S4). Furthermore, for each species, we selected the proteins detected in the three biological samples and classified them as HF core proteins (Table S5). The *E. granulosus* and *E. ortleppi* HF cores comprehended 61 and 105 proteins, respectively, with 52 being shared by the two species (Table 1).

**Table 1.**
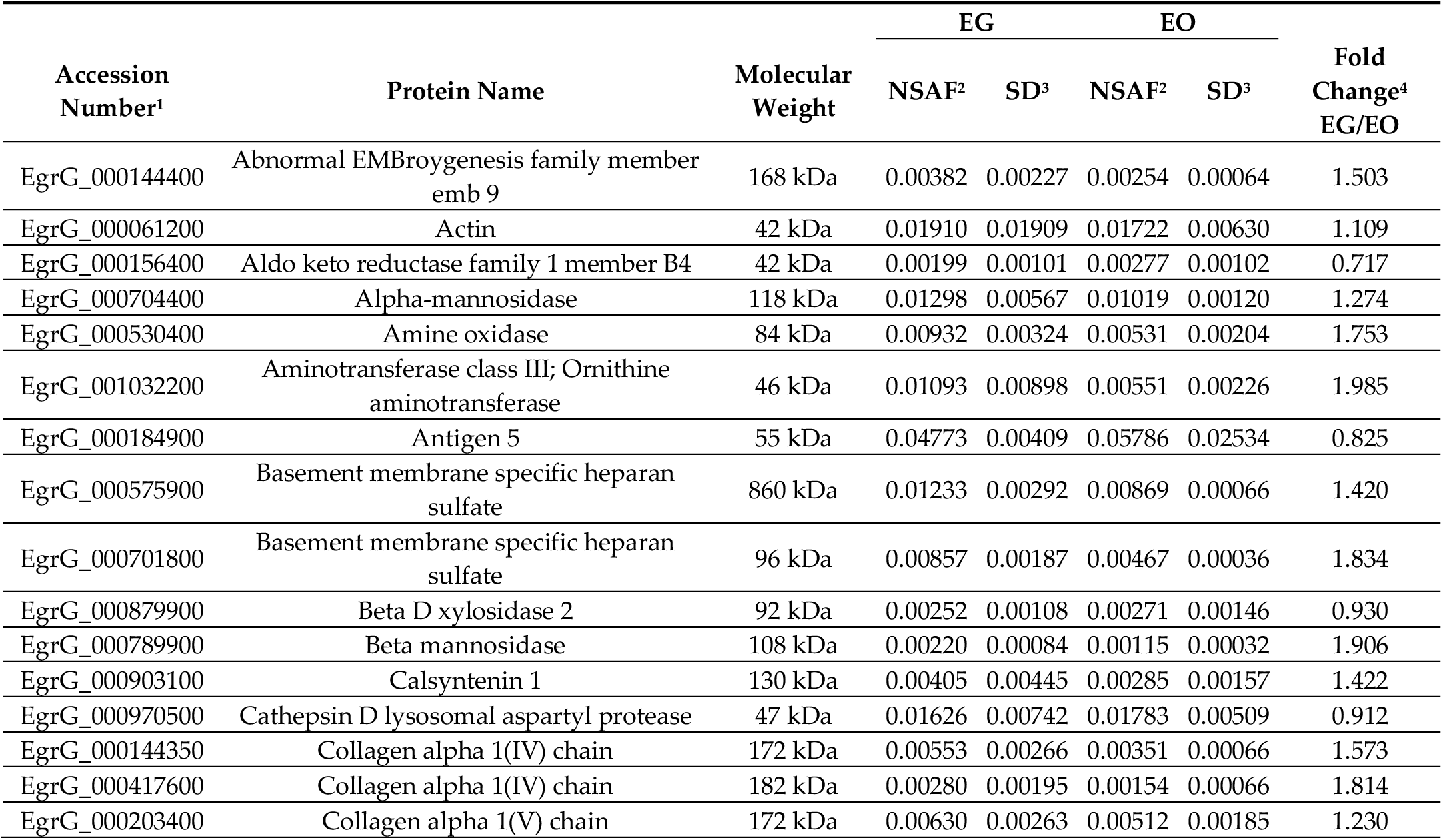
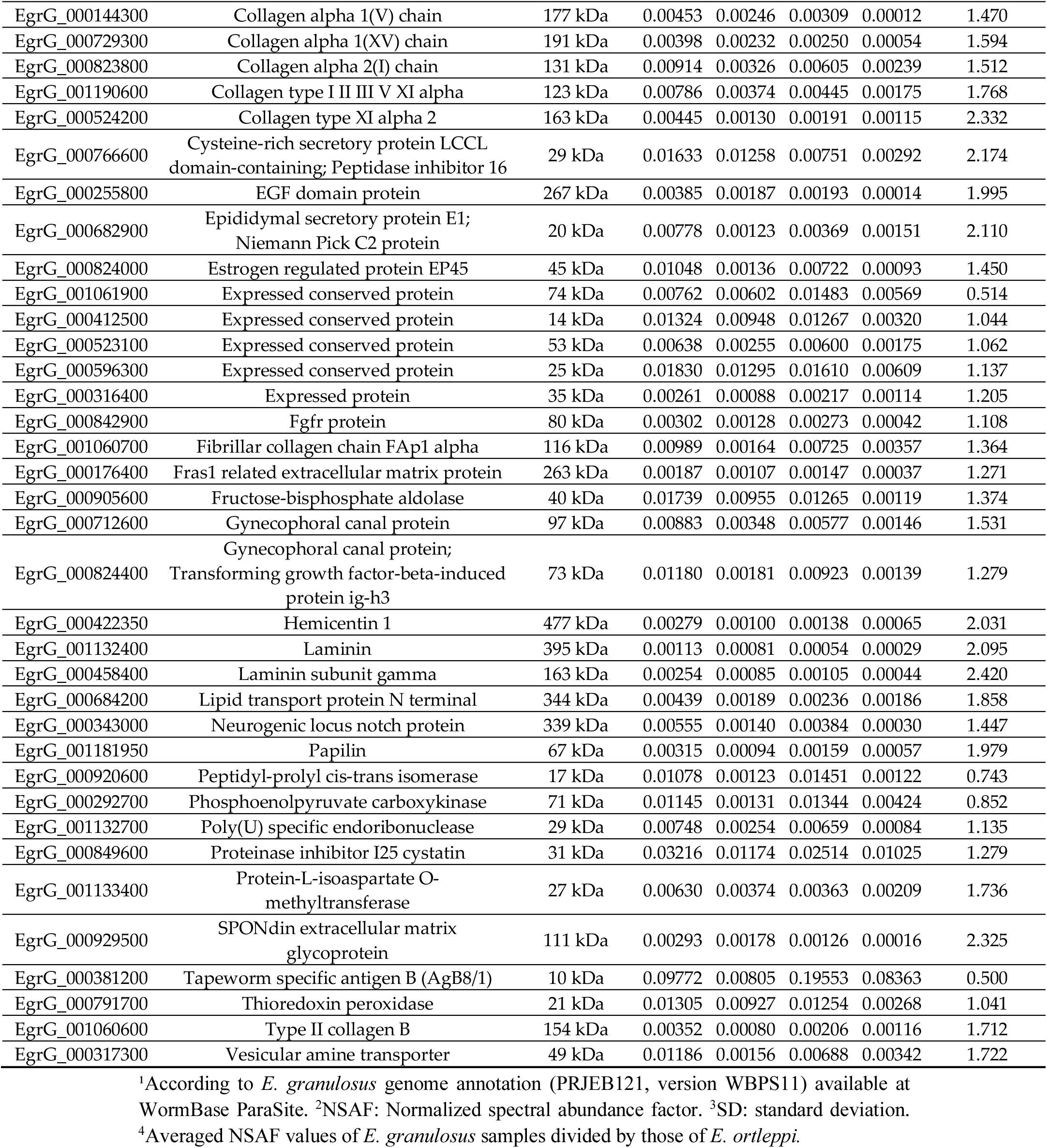
Core proteins identified in *E. granulosus* and *E. ortleppi* HF samples collected from pulmonary hydatid cysts. Listed proteins were identified in the three biological replicates from each species. Quantitative data are presented based on averaged NSAF values calculated for *E. granulosus* (EG) and *E. ortleppi* (EO).

In the shared core subgroup we identified proteins associated to different biological processes, such as cathepsin D, laminin, thioredoxin peroxidase, poly(U) endoribonuclease, cystatin, fructose-bisphosphate aldolase, and antigens previously described as relevant in *Echinococcus* spp. biology, such as antigen B (AgB) and antigen 5 (Ag5). AgB and Ag5 are antigens with recognized significance in *Echinococcus* spp. biology by their abundance and immunogenicity.

AgB is an oligomeric lipoprotein and its final structure can be composed by until five subunits (AgB8/1 to 5). We detected subunit AgB8/1 in the shared core subgroup of proteins, while subunits AgB8/2 to 5 were detected in only one *E. granulosus* sample (Figure S1 and Table S2). The levels of these subunits in the other samples might not be high enough to be detected under our experimental conditions.

The shared core proteins constitute the major proteins in the HF of *E. granulosus* and *E. ortleppi* and are interesting study targets to understand molecular mechanisms at the host-parasite interface in cystic echinococcosis, as well as, candidates to new therapies.

### 2.2. Potential Secretion Pathways Associated to the Parasitic Proteins Identified in E. granulosus and E. ortleppi Hydatid Fluid

The complete repertoires of *E. granulosus* and *E. ortleppi* proteins identified in the corresponding HF samples were analyzed by bioinformatic tools to predict whether identified proteins would be secreted by a classical pathway (signal peptide) or by an alternative pathway, and the results are summarized in Figure 2. In the *E. granulosus* repertoire (Figure 2a), 54% (150 out of 278) of the proteins were predicted to have a signal peptide, 11% (31 out of 278) were predicted as secreted by an alternative pathway and 35% (97 out of 278) were not predicted as secreted. In the *E. ortleppi* repertoire (Figure 2b), 45% (111 out of 249) of the proteins were predicted to have a signal peptide, 13% (32 out of 249) were predicted as secreted by an alternative pathway and 43% (106 out of 249) were not predicted as secreted.

**Figure 2.**
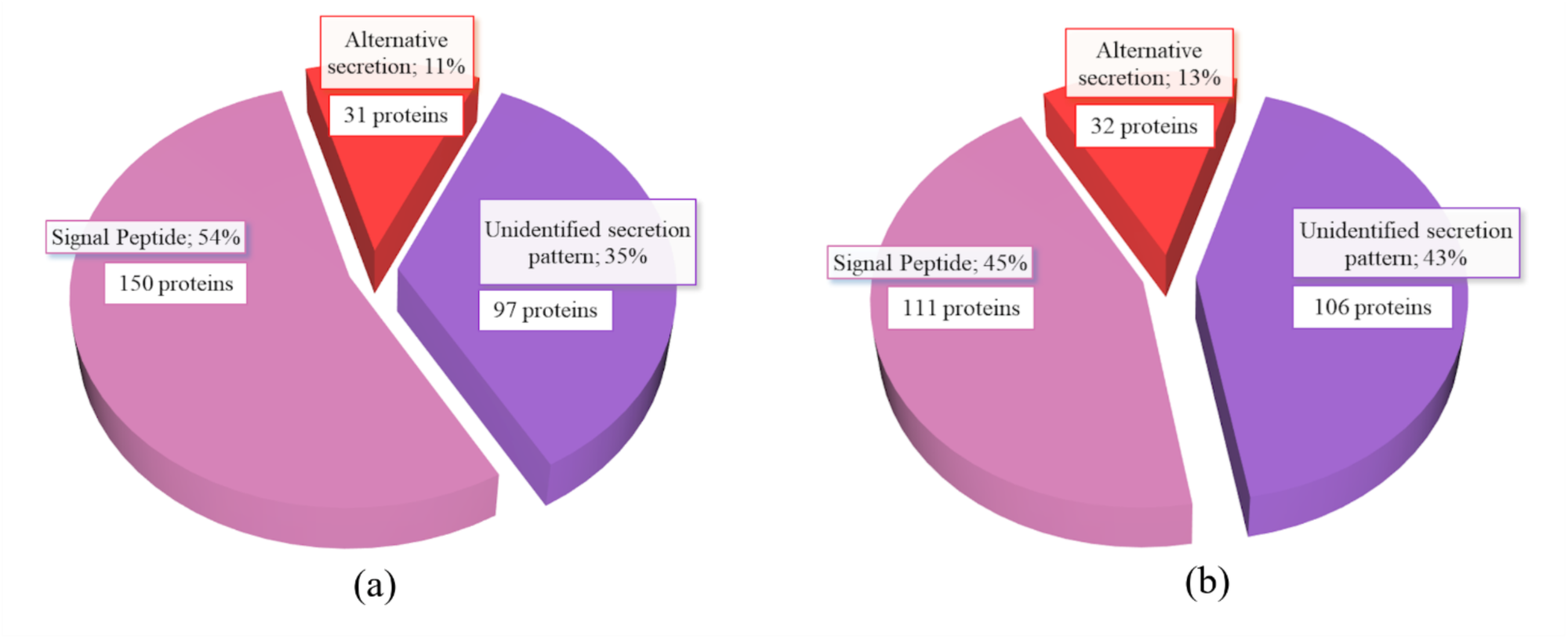
*In silico* prediction of secretion pathways for the *E. granulosus* and *E. ortleppi* repertoires of HF proteins. Percentages of the total and absolute number of proteins with probable classic or alternative signals for secretion are presented. Proteins with any identifiable signal for secretion were grouped under the term “Unidentified secretion pattern”. (a) *E. granulosus* proteins. (b) *E. ortleppi* proteins.

### 2.3. Insights Into the Molecular Mechanisms Associated to the Protein Repertoires of Hydatid Fluid from E. granulosus and E. ortleppi

Parasitic proteins detected exclusively in HF from *E. granulosus*, detected exclusively in HF from *E. ortleppi* or shared by HF samples from both species were classified according to COG (Clusters of Orthologous Groups) functional terms. Along the three groups, the most represented COG functional categories were “Posttranslational modification, protein turnover, chaperones” and “Carbohydrate transport and metabolism”, suggesting that proteins associated with these functions have major roles at host-parasite interface (Figure 3). The category “Signal transduction mechanisms” accounted for 17% (37 out of 218) of the sequences in the shared set of proteins, and the set of proteins detected exclusively in *E. granulosus* showed, additionally, 25.4% (18 out of 71) of the sequences associated to this category (Figure 3a and 3b). The category “Cytoskeleton” accounted for 5% (11 out of 218) of the proteins in the shared set, and the set of proteins detected exclusively in *E. ortleppi* showed a high proportion of sequences associated to this category (23.7%; 9 out of 38) (Fig 3a and 3c).

**Figure 3.**
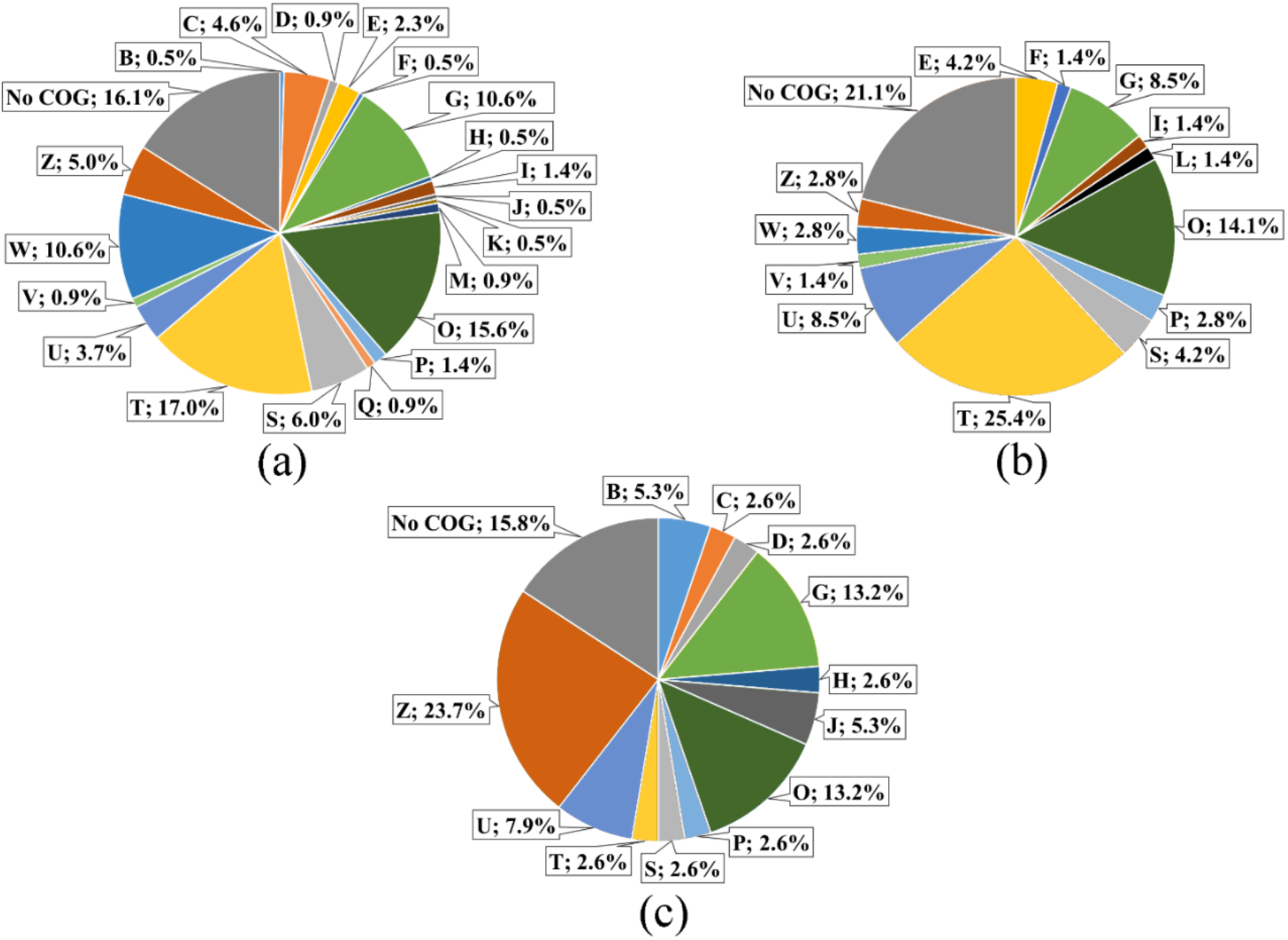
Functional analysis of parasitic proteins identified in HF samples from bovine pulmonary hydatid cysts. (a) Functional COG categories of parasitic proteins shared between *E. granulosus* and *E. ortleppi* samples. (b) Functional COG categories of parasitic proteins detected exclusively in *E. granulosus* samples. (c) Functional COG categories of parasitic proteins detected exclusively in *E. ortleppi* samples. Percentages of identified proteins in each functional category are indicated. B, Chromatin structure and dynamics; C, Energy production and conversion; D, Cell cycle control, cell division, chromosome partitioning; E, Amino acid transport and metabolism; F, Nucleotide transport and metabolism; G, Carbohydrate transport and metabolism; H, Coenzyme transport and metabolism; I, Lipid transport and metabolism; J, Translation, ribosomal structure and biogenesis; K, Transcription; L, Replication, recombination and repair; M, Cell wall/membrane/envelope biogenesis; O, Posttranslational modification, protein turnover, chaperones; P, Inorganic ion transport and metabolism; Q, Secondary metabolites biosynthesis, transport and catabolism; S, Function unknown; T, Signal transduction mechanisms; U, Intracellular trafficking, secretion, and vesicular transport; V, Defense mechanisms; W, Extracellular structures; Z, Cytoskeleton.

Host proteins identified in *E. granulosus* and *E. ortleppi* HF samples were also classified according to COG. The most represented COG categories in the group of shared host proteins were “Carbohydrate transport and metabolism” (6 out 25) and “Cytoskeleton” (7 out 25), followed by “Energy production and conversion” (3 out 25) and “Posttranslational modification, protein turnover, chaperones” (4 out 25) (Figure 4a). The latter was, additionally, well represented in the repertoires of bovine proteins identified exclusively in *E. granulosus* (4 out 12, Figure 4b) and *E. ortleppi* (4 out 16, Figure 4c). Thus, host proteins of different categories can penetrate into the hydatid cyst, but their functions there remain to be elucidated.

**Figure 4.**
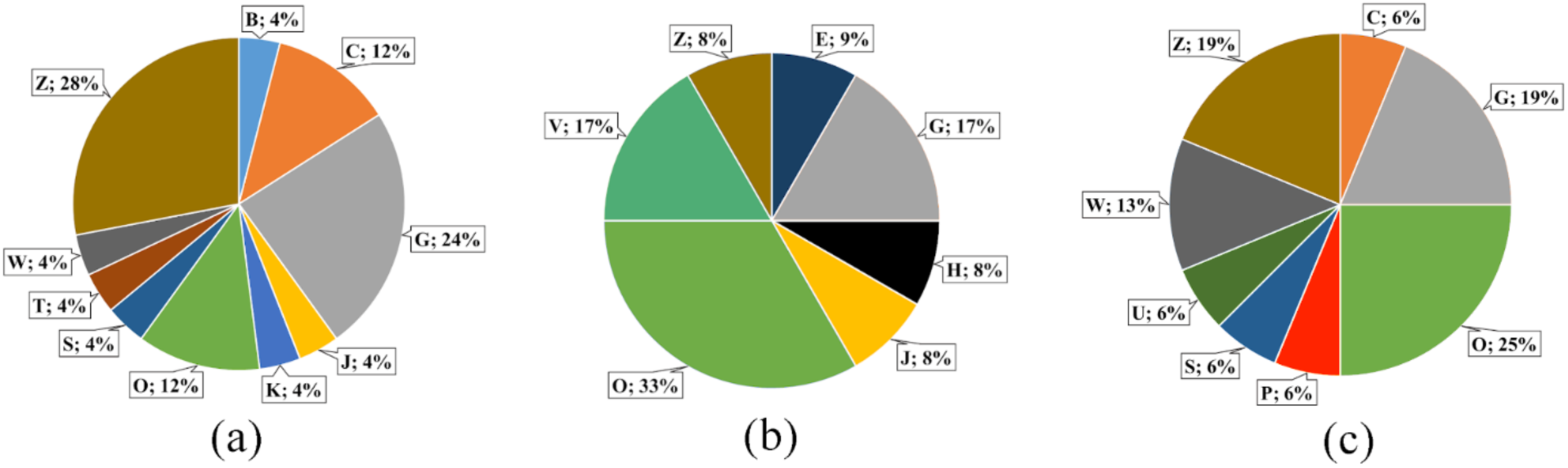
Functional analysis of host proteins identified in HF samples from bovine pulmonary hydatid cysts. (a) Functional COG categories of bovine proteins identified in *E. granulosus* and *E. ortleppi* samples. (b) Functional COG categories of bovine proteins identified exclusively in *E. granulosus* samples. (c) Functional COG categories of bovine proteins identified exclusively in *E. ortleppi* samples. Percentages of identified proteins in each functional category are indicated. B, Chromatin structure and dynamics; C, Energy production and conversion; E, Amino acid transport and metabolism; G, Carbohydrate transport and metabolism; H, Coenzyme transport and metabolism; J, Translation, ribosomal structure and biogenesis; K, Transcription; O, Posttranslational modification, protein turnover, chaperones; P, Inorganic ion transport and metabolism; S, Function unknown; T, Signal transduction mechanisms; U, Intracellular trafficking, secretion, and vesicular transport; V, Defense mechanisms; W, Extracellular structures; Z, Cytoskeleton.

Gene Ontology (GO) enrichment analyses were performed for the whole set of parasitic proteins identified in *E. granulosus* and *E. ortleppi* with the Cytoscape plugin BiNGO [23]. Functional classification with GO enrichment data is shown in supplementary Table 6. Most proteins were functionally annotated for both *E. granulosus* (220 out of 278 proteins) and *E. ortleppi* (203 out of 249 proteins). GO enrichment (p ≤ 0.05) was found for 180 GO subcategories in *E. granulosus* (Table S6A) and for 224 GO subcategories in *E. ortleppi* (Table S6B), considering the three main GO categories, biological process, molecular function and cellular component. *E. granulosus* and *E. ortleppi* showed the same profile regarding the most significant GO subcategories (p < 0.001), an additional indicative of the similar molecular strategies employed by these parasites at their larval stage.

The enriched GO terms, in biological process and molecular function major categories, for *E. granulosus* and *E. ortleppi* proteins were summarized by REVIGO [24]. The complete lists of summarized non-redundant terms are shown in supplementary tables 7 and 8. After the summarization by REVIGO, 65 and 63 category clusters were generated for *E. granulosus* and *E. ortleppi*, respectively. Considering the biological process major category, the clusters “cell adhesion”, “carbohydrate metabolic process” and “regulation of proteolysis” were among the most enriched clusters in both *E. granulosus* and *E. ortleppi* (Figures 5a and 6a). In the molecular function major category, the clusters “extracellular matrix structural constituent”, “calcium binding” and “hydrolase activity, acting on glycosyl bonds” were among the most enriched clusters in both *E. granulosus* and *E. ortleppi* (Figures 5b and 6b).

**Figure 5.**
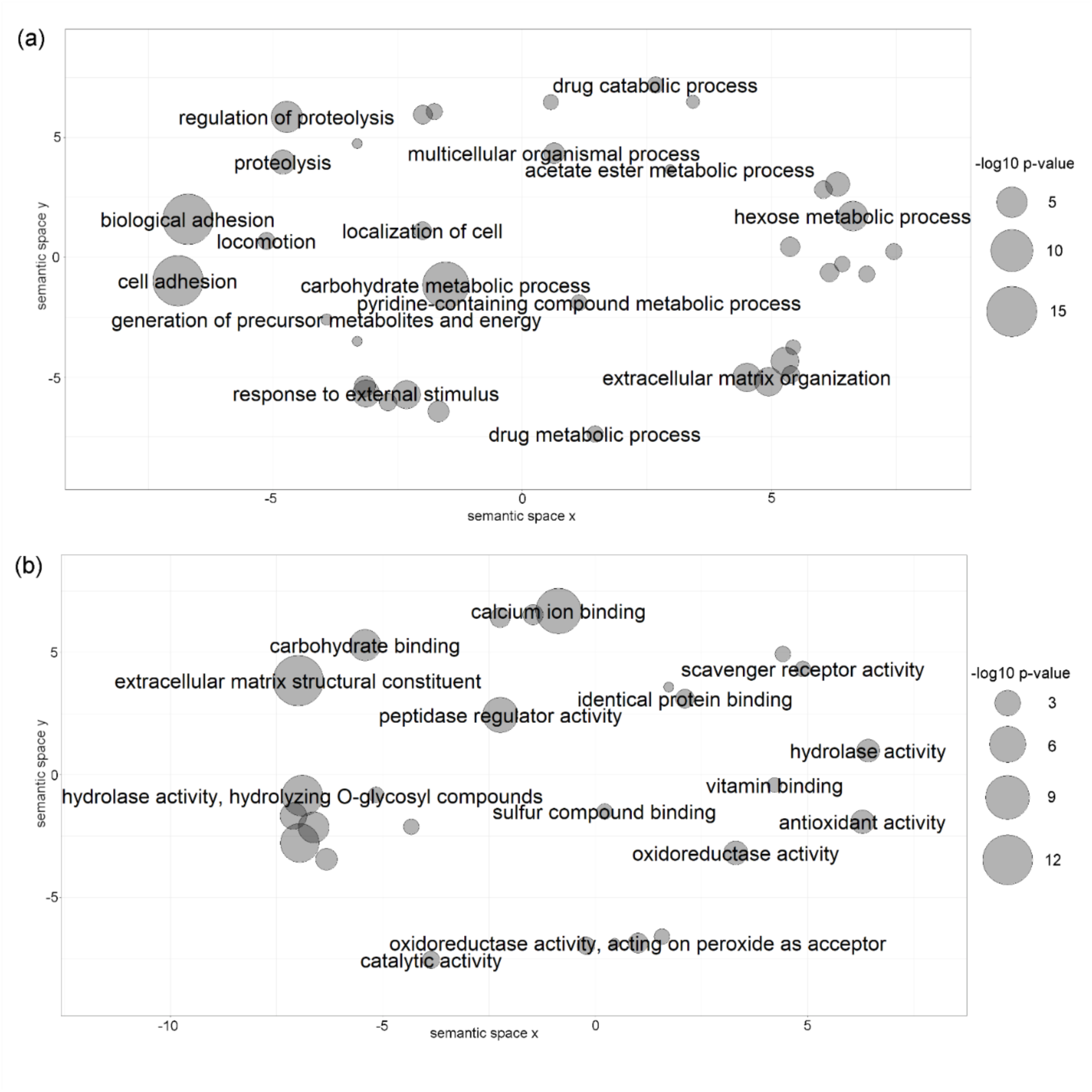
Category clusters of summarized related GO terms obtained in functional enrichment analysis of total set of identified proteins in *E. granulosus* HF. Scatterplot view of REVIGO clusters in major categories: (a) biological process; (b) molecular function. Sphere size is proportional to the p-value (larger spheres indicate more significant p-values, according to the scale).

**Figure 6.**
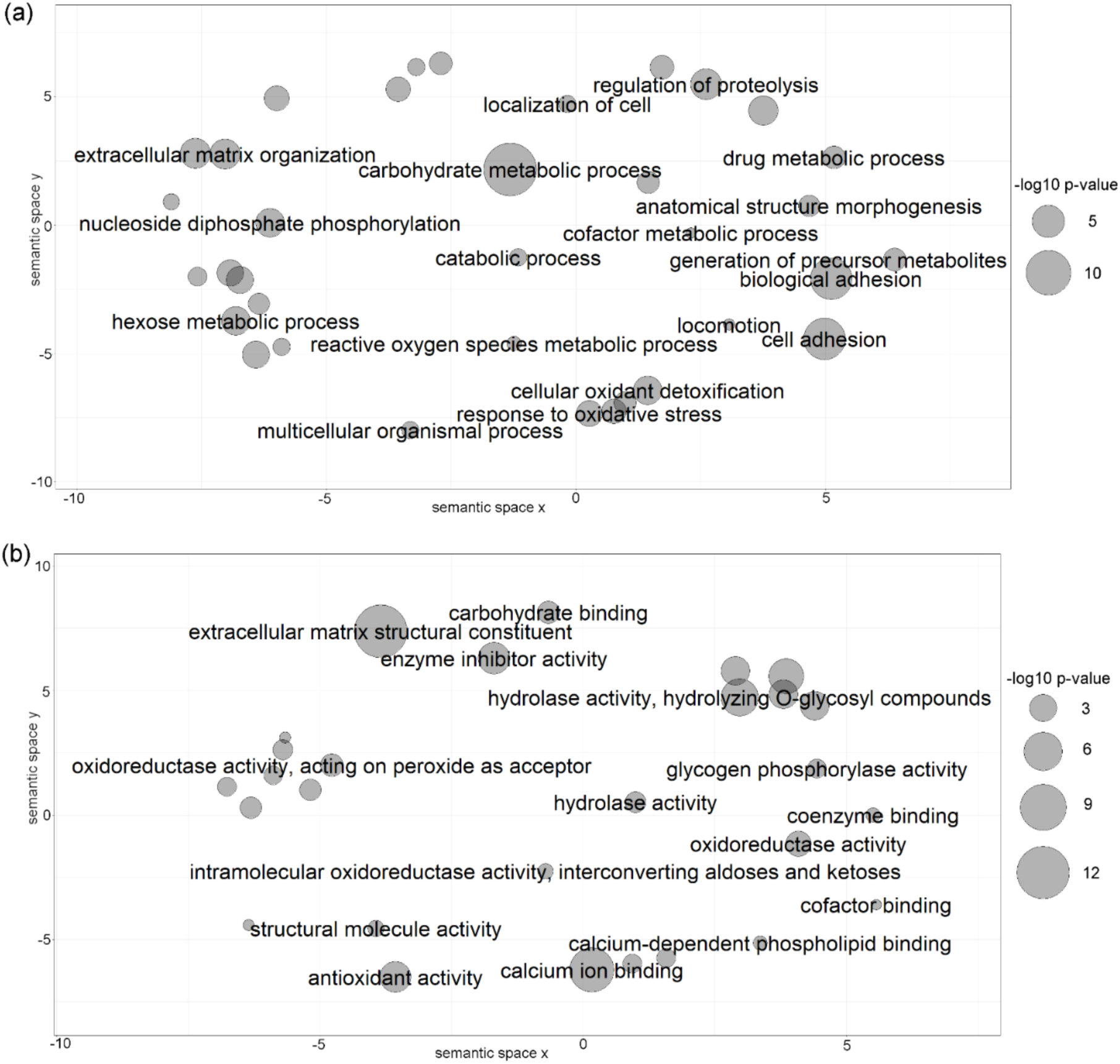
Category clusters of summarized related GO terms obtained in functional enrichment analysis of total set of identified proteins in *E. ortleppi* HF. Scatterplot view of REVIGO clusters in major categories: (a) biological process; (b) molecular function. Sphere size is proportional to the p-value (larger spheres indicate more significant p-values, according to the scale).

## 3. Discussion

In our study, we described the proteomic analysis and comparison of HF samples from three *E. granulosus* and three *E. ortleppi* hydatid cysts collected from *B. taurus* lungs. With the use of protein in-solution tryptic digestion, and peptide fractionation by SCX followed by LC-MS/MS, we identified totals of 280 and 251 proteins in *E. granulosus* and *E. ortleppi* samples, respectively, with an overall number of 317 different parasitic protein species. The comparison of the identified protein repertoires in the biological replicates allowed us to uncover a core of 52 proteins found in *E. granulosus* and *E. ortleppi* HFs. In this core protein set, we highlight the presence of proteins associated with extracellular matrix structures and dynamics, like collagen, laminin, hemicentin-1, SPONdin extracellular matrix glycoprotein, basement membrane specific heparan sulfate and FRAS1-related extracellular matrix protein 1. These proteins may be related to the maintenance of the structural integrity of the hydatid cyst wall, helping the metacestode resist to the host responses. Additionally, these proteins might modulate cell adhesion and matrix remodeling at host interface to facilitate cyst development. The mentioned proteins were identified in *E. granulosus* extracellular vesicles isolated from HF of sheep hydatid cysts [25]. The germinative layer, that forms the internal face of the cyst wall, is probably responsible for extracellular vesicles secretion. We hypothesized that production of extracellular vesicles containing those proteins could be a strategy to spread them to the entire cyst wall extension, as a form of coordinating processes at distinct positions in the germinative layer. The secretion activity of germinative layer occurs towards the inside, as well as, outward the hydatid cyst, so there is a possibility that these proteins could also act in the host tissue nearby.

Papilin was identified in the core repertoires of *E. granulosus* and *E. ortleppi* in this study. Papilin belongs to the ADAMTS (A Disintegrin-like and Metalloprotease with Thrombospondin Type 1 Motif) superfamily of proteins, more specifically to the ADAMTS-like subgroup, encompassing those proteins with high sequence homology to ADAMTSs, but lacking the catalytic domain [26]. Papilin is able to interact with extracellular matrix proteins and may modulate the activities of other ADAMTSs, especially inhibiting procollagen N-proteinase [27–30]. ADAMTS3, a procollagen N-proteinase, was also identified in the protein repertoires of *E. granulosus* and *E. ortleppi* and could be regulated by papilin. Additionally, a variety of collagen proteins were identified in the complete repertoires of *E. granulosus* and *E. ortleppi*, which could be the substrates for procollagen N-proteinases [26,31]. Besides ADAMTS3, other two proteins associated to collagen maturation and modification were identified, procollagen galactosyltransferase 2 and procollagen-lysine 2-oxoglutarate 5-dioxygenase [32,33].

Altogether, these findings could be associated to the cyst wall integrity and to its components, the germinative and laminated layers, functionality. The germinative layer inside the hydatid cyst has a pivotal role in hydatid cyst development and survival. The outward face of germinative layer is covered by a syncytial tegument that is also a physical barrier to the entrance of macromolecules into hydatid cysts [6,34]. The laminated layer, an acellular, carbohydrate-rich sheath secreted by the germinative layer, shields the parasite from direct attack by host immune cells [35]. The extracellular matrix proteins and their regulators may be associated to a molecular network that both keeps the integrity of the cyst wall and allows the tissue expansion necessary for hydatid growth.

Proteins associated to transport and metabolism of nutrients are well represented in our analysis. These proteins may act in basic cellular functions, playing important roles in nutrient uptake and production of structural constituents and energy. Some of them were found in the core protein repertoires of *E. granulosus* and *E. ortleppi* HF, like beta mannosidase, fructose-bisphosphate aldolase, phosphoenolpyruvate carboxykinase, aminotransferase class III, AgB and lipid transport protein N terminal. AgB is also the major antigen in HF and it has important immunomodulatory properties [7]. Of all five AgB subunits, we detected only AgB8/1 in all the six samples. AgB8/1 has been reported as the most abundant subunit in *E. granulosus* AgB oligomer [36], and it is the AgB subunit consistently identified by MS based analysis of HF [8,14,37]. However, the detection and the abundance of AgB8/2 to 5 in HF is rather variable [8,9,14,37]. Also, the AgB subunits representation in HF vary among different *Echinococcus* species [14,38].

Different carbohydrate-metabolizing enzymes were identified in the complete protein repertoires of HF from *E. granulosus* and *E. ortleppi*. Carbohydrate-metabolizing enzymes are repeatedly observed in the secretome of *E. granulosus* and other cestodes, including *E. multilocularis* [13,37,39]. Previous studies indicated that some carbohydrate-metabolizing enzymes exerted moonlighting effects along with their primary biochemical roles [40]. These molecules did not bear signal peptide, but significant amounts appeared to be secreted, probably through specific mechanisms, such as extracellular vesicles, to play a role extracellularly [37,41]. In addition to their primary described function, the carbohydrate-metabolizing enzymes identified in this study might exert moonlighting functions, protecting parasite’s tissues from host immune attack and aiding in metacestode development. Glycolytic enzymes have been found exerting a plethora of effects, such as binding to complement proteins and interference in their response, binding of host plasminogen and further increase in its activation, and interaction with adhesins and cytoskeleton to facilitate invasion [42–44]. In *E. granulosus*, it was demonstrated that fructose-bisphosphate aldolase interacts with actin [41].

Proteins acting in signaling pathways were identified in our analyses, many of them shared between *E. granulosus* and *E. ortleppi*. Desert hedgehog protein (Dhh), noggin, notch, tyrosine protein kinase otk and glypican-1 are examples of signaling proteins that play crucial roles in embrionary and morphological development in model organisms, such as *Caenorhabditis elegans, Drosophila melanogaster* and *Mus musculus* [45–48]. The germinative layer in fertile metacestodes is constituted of cells that actively participate in cyst development, these cells differentiate to generate brood capsules and PSC, secrete part of HF components and produce the molecules needed to maintain cyst wall integrity [6,49]. In this work, only viable fertile hydatid cysts were used, so the germinative layer was probably very active and the signaling proteins we found could have a function coordinating the events along the thin cell layer in the germinative tissue. As with the proteins associated with extracellular matrix, extracellular vesicles might carry these signaling molecules to different cyst compartments, allowing extracellular communication at the metacestode tissues.

Some proteins identified here are linked to the major developmental pathways, Hedgehog, Notch and Wnt, which are involved in many embryological development cascades, as well as cell fate, cell polarity and in maintaining stemness of stem cells [47,48,50]. Given that part of the cells in germinative layer are stem cells responsible for generating other cell types and tissues in the metacestode, our findings suggest that such developmental pathways are active in the *Echinococcus* spp. hydatid cyst. Differential expression of signaling proteins along different stages of development of *E. granulosus* and *Hymenolepis microstoma* has been recently demonstrated in the literature [51,52]. In *E. multilocularis* and *H. microstoma*, the expression patterns of Wnt proteins during larval metamorphosis have been elucidated [53]. The roles played by the signaling transducing proteins might be necessary for the proper development and growth of the metacestode.

*E. granulosus* and *E. ortleppi* HF showed a diverse range of proteolytic enzymes. Aminopeptidases, carboxypeptidases, cysteine peptidases, metallopeptidases, an enteropeptidase and others were identified in our proteomic analysis. Proteolytic enzymes have pivotal roles at host-parasite interface, especially related to nutrient acquisition, tissue migration and protection against host immune response [54–57]. Metalloproteases, another class of proteolytic enzymes frequently found in parasitic secretomes, function mainly in extracellular matrix degradation and tissue remodeling, but also facilitate a diverse range of cellular processes, including regulation of stem cell proliferation in planarians [58]. Based in their importance in different processes of basic parasitic biology and role at host-parasite interface, some proteases have been proposed as therapeutic targets [59–62].

On the other hand, protease inhibitors such as cystatins, serpins and proteins containing Kunitz and Kazal domains were also detected in *E. granulosus* and *E. ortleppi* HF. Four of these protease inhibitors were found in the core set of HF proteins. Proteases are part of defense mechanisms in mammals and the presence of parasitic protease inhibitors suggest that modulation of host protease activities could be a general mechanism of protection against elimination in *Echinococcus* spp. Furthermore, proteases and inhibitors could also be associated to the same molecular processes, where the inhibitors could be regulating protease activity to avoid excessive damage to the tissue [63]. In this case, the parasite would produce inhibitors to modulate the activity of their own proteases in order to keep the damage to the host tissue minimal, avoiding an increase in immune response at the infection site. Important immunomodulatory roles have been described for protease inhibitors in other parasitic flatworms [63,64]. Interestingly, in different invertebrates, Kunitz proteins have been described acting in defense against microbial infection and with toxin activity mediated by ion channels blockade [65,66].

Our comparative analysis showed a high degree of similarity between the protein repertoires of *E. granulosus* and *E. ortleppi* HF, which was expected considering the phylogenetic proximity of these two species. These results could aid in the prospection of new therapeutic interventions to cystic echinococcosis independently if caused by *E. granulosus* or *E. ortleppi*. Nevertheless, some proteins were identified exclusively in only one species, which could be indicative of some differential molecular mechanisms employed by each species to survive into the host organism. These proteins, identified exclusively in one species, could be explored as markers to determine the etiologic agent in cystic echinococcosis cases, and improve clinical end epidemiological data. However, it is important to notice that most of exclusive proteins were identified in just one biological replicate, which suggest that these proteins are necessary at specific infection contexts. They may represent molecular events triggered at particular time points in metacestode development, or yet to counteract a specific host response.

Looking at the set of exclusively identified proteins of each species, we found three cathepsin L cysteine proteases in *E. granulosus*. On the other hand, in *E. ortleppi* exclusively identified protein set, we found Calpain-A, a Ca^2+^-dependent cysteine protease with important roles in signal transduction, cytoskeleton rearrangement and microvesicles formation and budding [67,68]. Although belonging to different classes of cysteine proteases, cathepsin L and calpain-A, could have similar biological roles as parasite defense molecules in *E. granulosus* and *E. ortleppi* infection. Cathepsins are proteases widely described as molecular players in helminthic infections, suppressing host immune response at host-parasite interface [57]. Calpains are associated to cell degeneration, as studies have reported that under Ca^2+^ imbalances calpain become activated and mediates apoptosis and necrosis [69–71]. Thus, a role for Calpain-A as a defense molecule inducing cell death at infiltrating and adjacent host cells in *E. ortleppi* infection is possible.

Another proteins possibly involved in mediating apoptosis, the annexins, were also exclusively found in *E. ortleppi* HF samples. In *Taenia solium*, annexin B1 has been detected colocalizing with neutrophils and eosinophils in the surrounding host-derived inflammatory sites, and induced apoptosis in human eosinophils *in vitro* [72]. Thus, annexins could also function as defense molecules in *E. ortleppi* infection and the possibility of being able to induce apoptosis, as calpain-A, raise the question whether *E. ortleppi* would employ apoptosis to deal with the host in higher levels than *E. granulosus*. Cell death by apoptosis is an important phenomenon in flatworm infections, and occurs in both parasitic and host tissues. In *E. granulosus*, apoptosis is associated to metacestode infertility, by modulating the production of PSCs at the germinative layer of hydatid cysts [73]. In *S. mansoni*, the addition of soluble egg antigens to PBMC (peripheral blood mononuclear cells) cultures from patients induced apoptosis in T lymphocytes, supporting a role of these antigens in the modulation of the immune response mediated by apoptosis [74]. Evidence of a role for apoptosis in immunomodulation was also found in *E. multilocularis*, by showing that its metacestode ES products induce apoptosis in murine dendritic cells *in vitro* [75]. Further investigations will be necessary to determine the existence of differential patterns of apoptosis between *E. granulosus* and *E. ortleppi*.

We also searched for host proteins in the HF samples and identified them in lower amount than parasitic proteins, but with high diversity among the biological replicates from both *E. granulosus* and *E. ortleppi*. This indicate that different classes of host proteins can access the interior of the hydatid cyst, possibly performing a variety of functions, but these remain unknown. The balance parasitic/host protein content in HF has been associated to the fertility condition of *E. granulosus* hydatid cysts, where fertile ones have predominant protein content from the parasite, while infertile hydatid cysts have a higher quantity of proteins from host [37]. Samples collected in this study were from fertile hydatid cysts, so the low number of host proteins identified is in agreement with the literature. Infertile hydatid cysts may have a weakened wall, more susceptible to host protein entry. As mentioned before, we identified a large set of parasitic proteins related do extracellular matrix and structure maintenance, supporting the idea that in fertile hydatids the wall is an important structure acting as a barrier to protect the parasite.

Despite some protein detected exclusively in hydatid fluid from *E. granulosus* and *E. ortleppi*, the overall repertoire of proteins indicates a high similarity between them. Indeed, many proteins identified exclusively have redundant functions, or are associated to the same metabolic pathways, as shared ones. *E. ortleppi* genome has not been sequenced yet, so the protein identification was all based on a database of protein sequences deduced from the genome of *E. granulosus*. For this reason, some *E. ortleppi* proteins may have remained undetected in this study. It is likely that more differences will be detected between the HF proteomes from the two species once the *E. ortleppi* genome becomes available.

Nevertheless, our protein profiles evidenced important mechanisms related to basic cellular processes and functions, such as adhesion, extracellular structures organization, development regulation and enzyme activity acting in the host-parasite interface. Moreover, it suggests that *E. granulosus* and *E. ortleppi* employ very similar molecular tools in the bovine infections, and better ways to cope with cystic echinococcosis could be developed exploring these general features.

## 4. Materials and Methods

### 4.1. Biological Material

*E. granulosus* and *E. ortleppi* hydatid cysts were obtained from lungs of cattle slaughtered at a commercial abattoir in the metropolitan region of Porto Alegre, RS (Brazil). Animal slaughtering was conducted according to Brazilian laws and under supervision of the *Serviço de Inspeção Federal* (Brazilian Sanitary Authority) of the Brazilian *Ministério da Agricultura, Pecuária e Abastecimento. Echinococcus* spp contaminated viscera, identified during mandatory meat inspection, were donated by the abattoir for use in this work.

Lungs were dissected and HF was aseptically aspirated from the hydatid cysts. The HF recovered from individual cysts were centrifuged at 10,000 *x* g for 15 min at 4 °C to sediment PSCs and debris [8]. Only HF samples from fertile cysts, *i.e*., with viable PSCs, and with similar sizes (4-6 cm in diameter) were used in the study. The PSCs collected from each cyst were used for species identification, which was done by high-resolution melting (HRM) [76]. We selected three individual *E. granulosus* and three individual *E. ortleppi* HF samples (EG1-3 and EO1-3, respectively) for the proteomic analysis.

### 4.2. Sample Preparation and Mass Spectrometry (MS) Analysis

Protein concentration of each HF sample was determined using Qubit(tm) and was qualitatively evaluated by SDS-PAGE (12% SDS-polyacrylamide gel). Proteins were in-solution digested with trypsin and fractionated by strong cation exchange (SCX) [37]. To release the peptides, phosphate buffer 5 mM (pH 3.0) was added to the SCX columns with a salt gradient as follows: 75 mM KCl (fraction A), 125 mM KCl (fraction B), 200 mM KCl (fraction C), 300 mM KCl (fraction D) and 400 mM KCl (fraction E). Each fraction was lyophilized and stored at −80°C until liquid chromatography-tandem mass spectrometry (LC-MS/MS) analysis.

The five SCX resulting fractions from each one of the six biological samples were analyzed individually, totalizing 30 LC≤MS/MS runs. The mixture of tryptic peptides corresponding to each SCX fraction was automatically loaded onto a C18 Jupiter pre-column (Phenomenex; bead diameter 10 μm; 100 μm x 50 mm) by an Easy-nLCII nano HPLC system (Thermo Scientific). After loading the samples in solvent A (0.1% formic acid), the peptides were subjected to chromatographic separation in reverse phase using a C18 AQUA column (Phenomenex; beads diameter 5 μm; 75 μm x 100 mm). Both the pre-column and the analytical column were packed in house. The peptides were eluted on a gradient of 5 - 35% of solvent B (0.1% formic acid in acetonitrile) in 60 min; 35 - 85% B in 5 min; 85% B in 5 min; 85 - 5% B in 2 min and; 5% B in 13 min, under a flow of 200 nL/min. Spray voltage was set at 1.8 kV, 200 °C, and the mass spectrometer was operated in positive, data dependent mode, in which one full MS scan was acquired in the m/z range of 300–1800 followed by MS/MS acquisition using collisional induced dissociation (CID) of the ten most intense ions from the MS scan using isolation window width of 3 m/z. MS spectra were acquired in the Orbitrap analyzer at 30,000 resolution (at 400 m/z). Dynamic exclusion was defined by a list size of 500 and exclusion duration of 90 s at repetition intervals of 30 seconds. For the survey (MS) scan a target value (AGC) of 1,000,000 and maximum injection time of 100 ms were set whereas the target value for the fragment ion (MS/MS) spectra was set to 10,000 and maximum injection time of 100 ms. The lower threshold for targeting precursor ions in the MS scans was 200 counts per scan. The raw files (*.raw) from the MS and MS/MS spectra were converted to the extension *.mgf (mascot generic format) using the MSconvert software (available at http://proteowizard.sourceforge.net).

### 4.3. Database Search and MS Data Analysis

For protein identification, the generated LC-MS/MS data were used for searches in local databases containing the deduced amino acid sequences from the *E. granulosus* genome assembly (PRJEB121), version WBPS11, available at WormBase ParaSite (http://parasite.wormbase.org), and the *Bos taurus* protein sequences obtained from UniProt/Swiss-Prot (Proteome ID: UP000009136).

Mascot Search Engine v. 2.3.02 (Matrix Science) was used for peptide and protein identifications. The search parameters consisted of carbamidomethylation as a fixed modification, oxidation of methionine as a variable modification, two trypsin missed cleavage and a tolerance of 10 ppm for precursor and 1 Da for fragment ions. Ion type was set as monoisotopic, and peptide charges 2+, 3+ and 4+ were taken into account.

Peptide and protein identifications were validated using Scaffold v. 4.8.7 (Proteome Software Inc.). The peptide identifications were accepted if they could be established with >95% probability. Protein identifications were accepted if they could be established at greater than 99% probability and contained two unique identified peptides. The false discovery rate (FDR) was 0.9% and 0.0% for proteins and peptides, respectively. The mass spectrometry proteomics data have been deposited to the ProteomeXchange Consortium via the PRIDE [77] partner repository with the dataset identifier PXD019314 and 10.6019/PXD019314.

Some histones (proteins known to be highly conserved in eukaryotes) did not fulfill the criteria of at least two unique peptides when the identifications obtained using each database, *E. granulosus* or *B. taurus*, were compared (Table S9). Because we were unable to assure their organism of origin, histones H4 (EgrG_000323100 and E1BBP7), H2A (EgrG_002051500 and A0A0A0MP90) and H2B (E1BGW2) were removed from further analysis.

Normalized spectral abundance factor (NSAF), acquired using Scaffold, was used to quantify the differences in protein abundance between samples [78]. To determine statistical differences between NSAF values of *E. granulosus* and *E. ortleppi* shared proteins, we performed a Students *t-*test and p-value correction with the Benjamini & Hochberg FDR. The heat map was performed in Matrix2png web interface (https://matrix2png.msl.ubc.ca/) with NSAF values of all identified proteins.

### 4.4. Prediction of Secretion Pathways

The identified parasite proteins were searched for the presence of a secretion signal peptide using SignalP 4.1, PrediSi and SecretomeP 2.0. The presence of an alternative signal for exportation was verified using SecretomeP 2.0. A protein was considered to contain a classical signal peptide when two out of three software detected a signal peptide sequence. Proteins that did not fit this criterion, but showed a NN score higher than 0.6 in Secretome P, were considered alternatively secreted proteins. Those that did not fulfilled any of the previous parameters composed the group of proteins with unidentified secretion pattern.

### 4.5. Functional Annotation

The EggNOG database (version 4.5.1, available at http://eggnogdb.embl.de/#/app/home) [79] was used to obtain the functional annotation of identified proteins from both parasitic and host origins. The functional annotation was performed separately to each set of identified protein: shared proteins (found in HF samples of *E. granulosus* and *E. ortleppi*), EG exclusively identified (found in HF samples of *E. granulosus* only) and EO exclusively identified (found in HF samples of *E. ortleppi* only).

Parasitic proteins were subjected to Gene Ontology (GO) enrichment analysis. The analysis was performed for the total repertoire of proteins from each species, using the Cytoscape plugin BiNGO [23]. The ontology files were retrieved from GO database, while Wellcome Trust Sanger Institute (UK) kindly provided the files associated with *E. granulosus* protein annotation. Functional enrichment analyses were performed using hypergeometric distribution and p-value correction with Benjamini & Hochberg FDR. Values of p ≤ 0.05 were considered statistically significant.

The platform REVIGO (http://revigo.irb.hr/) was used to remove redundant GO terms and summarize the lists of enriched GO terms [24]. For this, the semantic similarity of the GO terms was calculated through SimRel (allowed similarity = 0.5).

## 5. Conclusions

Our proteomic analysis highlighted molecular mechanisms operating at the host-parasite interface during *E. granulosus* and *E. ortleppi* infections in lungs from bovine hosts. The presence of proteins related to extracellular matrix organization and dynamics, developmental pathways, signaling transduction and carbohydrate metabolism in the HF was remarkable. The results provide valuable information on mechanisms shared by or differential between *E. granulosus* and *E. ortleppi*, helping to understand biological aspects of cystic echinococcosis caused by different parasite species. Moreover, they contribute to the knowledge on *E. ortleppi*, a species still poorly characterized molecularly. The *E. granulosus* and *E. ortleppi* protein profiles generated in our work can guide the choice of specific molecular processes to be analyzed in new comparative studies of these two species. Some of the identified proteins can be of clinical and epidemiological interest, because they can be used to more easily distinguish between human cystic echinococcosis caused by *E. granulosus* or *E. ortleppi*. Others and the pathways they belong to can be exploited to develop novel and more effective therapies against these two and other *Echinococcus* species.

## Supplementary Materials

Figure S1: Heat map of parasitic proteins identified in HF samples. All identified proteins are represented (blue: lower abundances; red: higher abundances), and their annotations are shown on the left, Figure S2: Bovine proteins identified in hydatid fluid samples from pulmonary cystic echinococcosis, Table S1: Parasitic proteins identified by LC-MS in *E. granulosus* and *E. ortleppi* samples, Table S2: Comparative analysis of proteins identified by LC-MS in *E. granulosus* and *E. ortleppi* samples, Table S3: Bovine proteins identified by LC-MS in *E. granulosus* and *E. ortleppi* samples, Table S4: Comparative analysis of the proteins detected in at least two biological replicates in one of the species, Table S5: Core proteins from *E. granulosus* and *E. ortleppi*, Table S6: Functional classification and gene ontology (GO) enrichment analysis of proteins detected in hydatid fluid of *E. granulosus* and *E. ortleppi*, Table S7: Summarized GO categorization of proteins detected in *E. granulosus* hydatid fluid, Table S8: Summarized GO categorization of proteins detected in *E. ortleppi* hydatid fluid. Table S9: Histone peptides identified in the LC-MS analysis using *E. granulosus* and *B. taurus* database,

## Author Contributions

Conceptualization, G.B.S., H.B.F. and A.Z.; methodology, G.B.S., K.M.M., J.C.L. and E.S.K.; formal analysis, E.D.S and J.C.L.; investigation, G.B.S., E.D.S., M.E.B. and E.S.K.; visualization, E.D.S., writing—original draft preparation, G.B.S., E.D.S. and A.Z.; writing—review and editing, E.D.S., K.M.M., J.C.L., E.S.K., H.B.F., S.M.T.S and A.Z.; supervision, A.Z.; project administration, A.Z.; funding acquisition, H.B.F., S.M.T.S. and A.Z. All authors have read and agreed to the published version of the manuscript.

