## Supplementary figures and images for "The secreted protein signature of hydatid fluid from pulmonary cystic echinococcosis"

### Figure S1

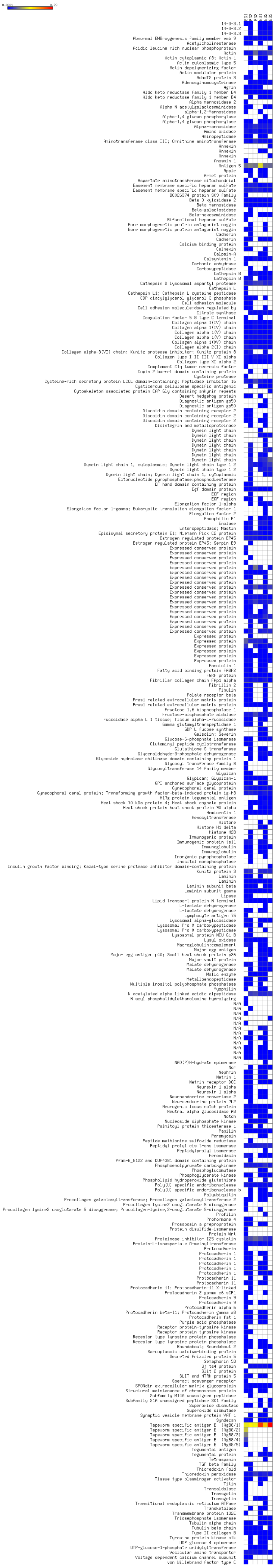

### Figure S2

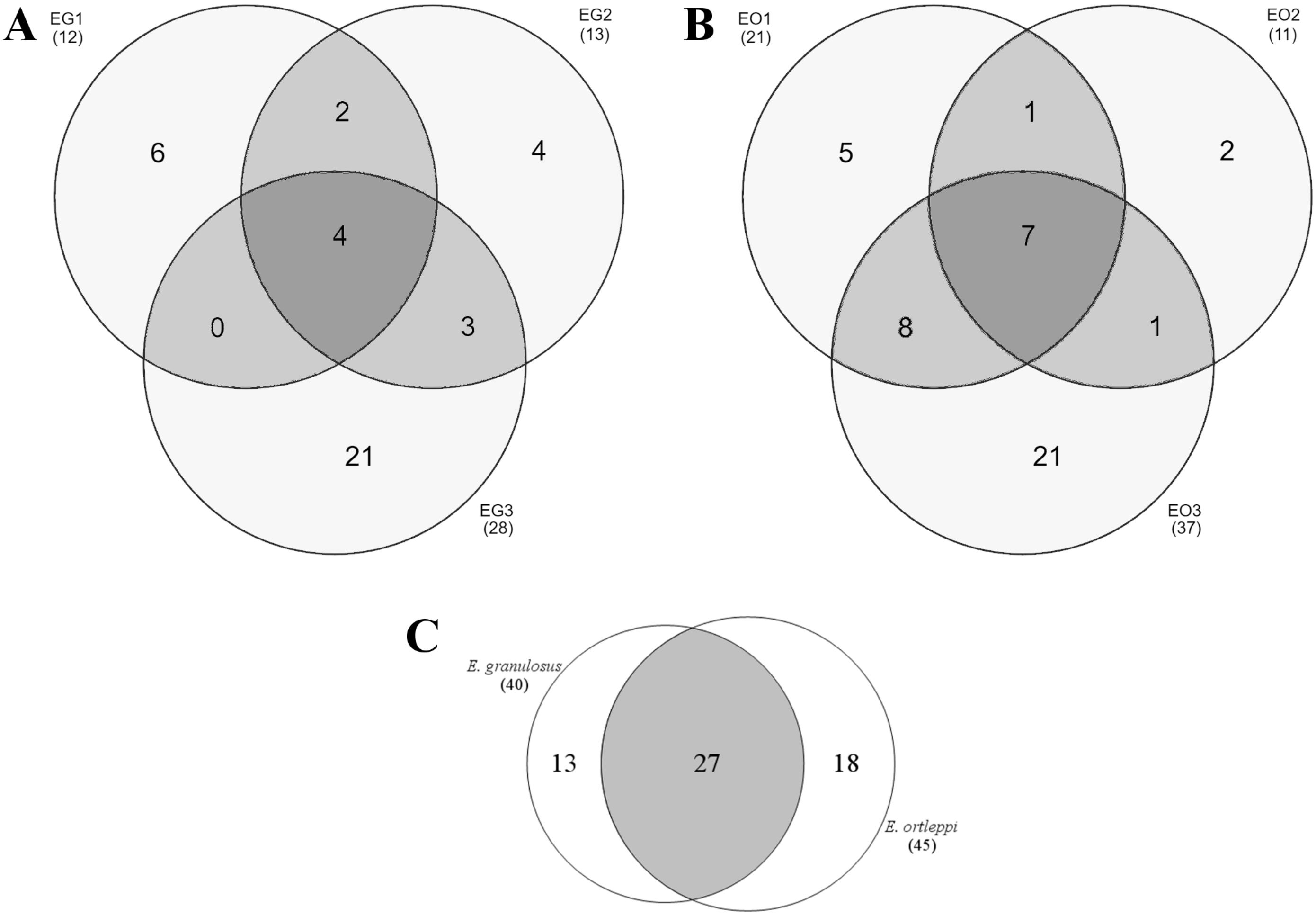
